# Source-sink dynamics explains the co-existence of the invasive pest *Dryocosmus kuriphilus* and its biological control agent *Torymus sinensis* across French Eastern Pyrenees

**DOI:** 10.1101/2025.03.27.645715

**Authors:** Jean-Loup Zitoun, Raphaël Rousseau, Sébastien Gourbière

## Abstract

Biological invasions have become a major cause of ecological and economic costs in many (agro-)ecosystems. Understanding the regulation of their local dynamics by resources limitation and natural enemies, i.e. ‘bottom-up’ and ‘top-down’ determinants, through inherently heterogenous environments stands as a critical challenge to provide efficient risk assessments and management measures. Here, we propose an unprecedented integrative modelling approach based on a spatial extension of the seminal Nicholson-Bailey model parameterized by a comprehensive field assessment of all the ‘bottom-up’ and ‘top-down’ determinants affecting the spread of a worldwide invasive pest, *Dryocosmus kuriphilus*, in 23 natural chestnut tree populations of the French Eastern Pyrenees. The analysis of the within-site dynamics allowed to quantify the spatial heterogeneity in the two types of regulatory forces across the study area, and to identify 16/23 sites where the control agent, *Torymus sinensis*, is expected to persist and control the invasive pest. The comparison of these predictions with the levels of *D. kuriphilus* hyperparasitism by *T. sinensis*, observed in all 23 forest sites, suggested hidden source-sink dynamics within the study area. The investigation of such dynamics by the coupling of any two local dynamics predicted in each of the 23 sites showed that i) low rates of *D. kuriphilus* and *T. sinensis* dispersal lead to a synchronization to the dynamic expected in the site with the highest chestnut tree frequency, while ii) higher rates typically allow for a stable equilibrium. Those predictions provide quantitative evidences for the persistence of *T. sinensis* in the 7 predicted ‘sink’ sites sustained by its local 16 ‘source’ populations. Dispersal then tends to homogenize the local efficiencies of the control agent while decreasing its global impact on the pest invasion. Overall, this pioneer spatial modelling of *D. kuriphilus* - *T. sinensis* interaction suggests that both introduced species are likely to persist in the European forest environments in a ‘co-invasion’ scenario.

## Introduction

Biological invasions have been steadily increasing in frequency since the beginning of the 19^th^ century and the rise of international trades (1,2). The increase in shipping rate over the last two centuries (3) has undoubtedly facilitated the spread of species outside of their natural ranges (4) where they rely on local resources to establish and grow as invasive populations (5). For herbivore insects, one of the most successful invasive taxa on all continents (6–8), crops and planted forests constitute key resources as they usually are made of monospecific groups with low genetic diversity that are associated with reduced abundances of predators and competitors (9,10). The damages caused by invasive insect pests to those valuable plant resources and their management costs induce strong economic losses that have reached up to US$26.8 billion a year over the past few decades (11). Beyond those impacts on field crops (12) and agroforestry (13), invasive herbivore insects represent one of the most important threats to ecosystems and biodiversity conservation (14,15). Given the broad impacts that invasive insect pests have on natural and agro-ecosystems and the services they provide (12, 16, 17), it has become critical to identify the key determinants of the local spread of invasive pests (18,19) to strengthen our ability to assess the corresponding risks and provide quantitative predictions on the potential efficacy of protection and mitigation measures (20,21).

While trying to establish into a local ecological network, invasive species face so-called ‘bottom-up’ and ‘top-down’ regulatory forces that effectively shape their population dynamics as they are associated with resources limitation and natural enemies, respectively (e.g. 22,23). The abundance, distribution, genetic and nutritional characteristics of host plants have all been shown to be ‘bottom-up’ determinants influencing the spread of invasive herbivore insects (18,24,25). Meanwhile, such invasive species represent ecological opportunities for native insect predators and parasitoids that can shift to those new hosts as they become more abundant (26,27). Although the inherent lack of co-evolutionary history can limit the efficiency of such native natural enemies (28), there are clear evidences that host shifts can contribute to the ‘top-down’ regulation of the invasion of insect pests (11). Last but not least, the introduction of parasitoids that evolved with the invasive species in their natural distribution area and the potential use of ‘improved’ native parasitoids, e.g. with increased levels of performance on the novel hosts, are key strategies for the management of invasive pests in emerging agricultural and agroforestry sustainable practices (29,30). While field evidences of the effects of the above bottom-up and top-down determinants have been essential to our understanding of the population ecology of herbivorous insect (see 31 for a review), their concomitant assessement through integrative tri-trophic modelling approaches are conspicuoulsy lacking, so that we have little quantitative knowledge about their relative importance in shaping the local dynamic of herbivore insects invasions (e.g. 18,32). The implementation of dynamical models bringing together field measures of those determinants therefore stands as a key challenge to better understand their interplay during invasive insect pest emergence and to anticipate the efficacy of biological control agents to protect crops, natural and planted forests (33,34).

The worldwide spread of the hymenopteran *Dryocosmus kuriphilus* (Yasumatsu, 1951) is a typical example of insect invasion, tightly associated with trade transports and representing a significant threat to many agro-ecosystems, and whose local emergence is affected by bottom-up and top-down determinants that include a parasitoid, globally used as biological control agent, *Torymus sinensis* (Kamijo, 1982). Within the last 50 years, this invasive gall forming parasite has spread from China to infest both cultivated and wild chestnut tree populations (*Castanea sativa* Mill., 1768) in Asia (35), the US (36) and Europe (37), causing up to 80% losses in chestnuts production (38), lowering tree biomass (39), and amplifying the impact of chestnut blight disease in natural forests (40). The availability of the host plant has been shown to facilitate the local spread of *D. kuriphilus* with chestnut tree dominated forests and monocultures suffering the highest rates of tree infestation (24,41) and pest-induced levels of defoliation (42). Meanwhile, genotype-dependent variation in susceptibility (43,44) also contribute explaining the variations in infestation observed between European and Asian *Castanea* species and their hybrids (45), between local ecotypes in Southern Italy (46), and at the individual tree level within populations of *C. sativa* (24,47). Along with these bottom-up determinants, *D. kuriphilus* natural enemies can contribute to its top-down regulation. In various places across Asia, North-America and Europe, native fungal (48,49) and hymenopteran (50–52) species have been shown to hyperparasitize its larvae inside the developping galls. While the measured rates of hyperparasitism by those native species remain lower than 10%, the chinese parasitoid, *T. sinensis*, that has been released in Japan (53), North-America (54) and Europe (55), typically infest over 75% of *D. kuriphilus* larvae (55,56), providing an efficient bio-control strategy (52).

In a recent eco-genomic study, we designed the first *C. sativa* - *D. kuriphilus* - *T. sinensis* dynamical model integrating estimates of all the above bottom-up and top-down determinants that we derived from a two years field study set in the French Eastern Pyrenees (24). This well empirically informed modelling showed that the emergence of *D. kuriphilus* in the chestnut tree forests has been predominantly shaped by the frequency and genetic susceptibility of chestnut trees until the introduction of *T. sinensis*. It further predicted that the initial reduction of the pest abundance by the control agent, which we observed in the field, is likely to be deceptive as it shall not allow for the co-extinction of the two exotic species, but for their long term coexistence with periodical re-emergences of the parasite as already described in Japan (57). While this integrative modelling provided clear quantitative insights into the relative importance of key bottom-up and top-down determinants in shaping the spread of this insect pest invasion, it did not allow to investigate the impact of their spatial variations on the dynamic of *D. kuriphilus* emergence and its interaction with the control agent *T. sinensis*.

In this contribution, we aim at expanding our understanding of the *C. sativa - D. kuriphilus - T. sinensis* interaction by considering its dynamics in heterogeneous forest environments where sites are characterized by different levels of bottom-up and top-down forces and connected by dispersal of the pest and its control agent. We first refined our comprehension of the within site tritrophic dynamic of interaction by analysing the effect of the absolute density of forest trees, and by prediting the spreading potential of *D. kuriphilus* and its control agent, *T. sinensis*, in each of the 23 forest study sites of the French Eastern Pyrenees where all the bottom-up and top-down determinants mentioned above were previously measured (24). The integration of the observed levels of heterogeneities of these determinants led to identify study sites corresponding to sources and sinks (58,59) for the invasive species and its control agent. The comparison of these predictions with the levels of chestnut tree infestation by *D. kuriphilus* and of *T. sinensis* hyperparasitism that we concomitantly characterized in the field (24), pointed toward hidden source-sink dynamics (60) within the study area. We then expanded our modelling into a two-site model to investigate the global dynamics resulting from the coupling of sites with local dynamics corresponding to those predicted in our 23 forest sites. This integrative eco-genomic modelling study provides new quantitative insights into the importance of *D. kuriphilus* and *T. sinensis* dispersal on the outcomes of the spatial interaction that contribute deciphering the local spread and biological control of this worldwide invasive pest.

## Materials and methods

### The C. sativa - D. kuriphilus - T. sinensis interactions in the French Eastern Pyrenees

The worldwide invasive gall forming hymenopteran *D. kuriphilus* is the most virulent pest of sweet chestnut tree (*C. sativa*) cultivated and wild populations (61). The chestnut tree populations located in the French Eastern Pyrenees predominantly correspond to chestnut coppices and patches of deciduous and coniferous forests where the tree density and frequency of *C. sativa* average ∼989 trees per hectare and ∼60%, with ranges of variation (between study sites of 1 hectare) spanning from 250 to 1570 and 15.8% to 90.8%, respectively (24). The insect pest has broadly spread into those chestnut tree forests according to a high intrinsic growth rate (*R_0_*) that was estimated to reach ∼15.9 (24). Such a strong invasive potential is commonly attributed to a highly effective parthenogenetic reproduction and a univoltine life-cycle timely synchronized with the chestnut tree host phenology (52). Semelparous females emerging from galls in June-July disperse during a 1-7 days long adult life-span to lay asexually produced eggs in chestnut tree buds (62). Their poor ability to navigate towards host plants during this short time period (63) makes such dispersal a merely diffusive process and their rate of oviposition to increase with the frequency of chestnut trees in the forest site (24,41). Oviposition triggers a local hypersensitive immune response in chestnut trees buds that can prevent the development of deposited eggs (46). Such host resistance varies between chestnut tree species (45) and local ecotypes (46) and was shown to allow for ∼68% of deposited eggs to survive in the studied chestnut tree forests (24). The *D. kuriphilus* eggs that escape this immune response and survive to intrinsic developmental failures, hatch within a month to develop into first instar larvae that enter in dormancy to overwinter. In spring, the development into second larval instars induces the formation of galls during the chestnut tree buds burst. *D. kuriphilus* individuals then become pupae and adults that will eventually emerge and produce the next and non-overlapping generation of eggs. Meanwhile, in invaded Asian, European and North-American chestnut tree populations, *D. kuriphilus* galls have been repeatedly shown to be hyper-parasitized by native fungi and hymenopteran parasitoid species (48–52). Such hyper-parasites were also observed in the French Eastern Pyrenees as *D. kuriphilus* larvae were found parasitized by endemic hymenopteran and fungal hyperparasitic species at rates reaching 4.5% and 5.4% on average, with variations between forest sites ranging from 0% to 27.3% and 0% to 17.7% (24). Those low values were all very consistent with previous field estimates obtained in neighbouring areas in south of France (51) and elsewhere in Europe (49,52). While such rates of hyperparasitism by native species were shown to play a secondary role in reducing *D. kuriphilus R_0_*, the studied chestnut tree populations were also heavily infested by the control agent, *Torymus sinensis*, with an average rate of infestation of 91% and spatial variations spanning between 57% and 100% across the entire area (24). Again, such prevalences of infestation were very similar to those previously reported in Europe (55,56).

### The tri-trophic model of interactions in a spatially heterogeneous environment

In order to investigate the dynamic of *C. sativa - D. kuriphilus - T. sinensis* interactions in an heterogeneous forest environment, we expanded our previous tri-trophic modelling (24) into a two-sites model. The dynamic of the tri-trophic interactions within each site was then described according to the model developed by Zitoun et al. (24), and the two sites were connected by the dispersal of the pest and its control agent.

*The within site dynamic of tri-trophic interactions.* The model of *C. sativa - D. kuriphilus - T. sinensis* interactions (24) was derived from the seminal host-parasitoid model proposed by Nicholson-Bailey (64) and adapted to account for the effects of the bottom-up and top-down factors regulating *D. kuriphilus* invasion. The Nicholson-Bailey framework is built on a description of the entangled host and parasitoid life cycles, which we tailored to represent the interaction between *D. kuriphilus* and its control agent *T. sinensis* (Figure 1). In late spring, the larvae of *D. kuriphilus* that have successfully developed inside the galls, referred to as H(t), can escape parasitism by *T. sinensis* adult females, denoted P(t), with probability F_s_. Those non-parasitized larvae can develop into pupae if they further escape native hymenopteran and fungal hyperparasites, according to probabilities F_ni_ and F_nf_, and survive to intrinsic causes of mortality at rate S_Hl_. The pupae then moult into adults with probability S_Ha_ and the emerging individuals in June-July disperse to lay their eggs. While each adult produce on average F_H_ eggs, only a fraction d of them are ultimately deposited in chestnut tree buds which, given the low ability of *D. kuriphilus* to navigate towards its host plant (63), was shown to be equal to the frequency p_c_ of chestnut tree in the local forest environment (24). Deposited eggs surviving the immune response of the trees and developmental failures, according to probabilities S_e_ and S_He_, hatch into first instar larvae that enter in overwintering dormancy. Finally, after the induction of gall development early in the spring of the following year, chambers formation leads to a typical ‘contest’ competition for space (65) resulting in a density dependent survival D of larvae to produce the new generation of successfully developed *D. kuriphillus* individuals in the galls. Those subsequently face the challenge set up by the new generation of *T. sinensis* adult females produced by the development at a rate S_P_ of *T. sinensis* larvae born from the eggs deposited in the larvae of the previous generation of *D. kuriphilus* infected with probability 1-Fs.

**Figure 1.**
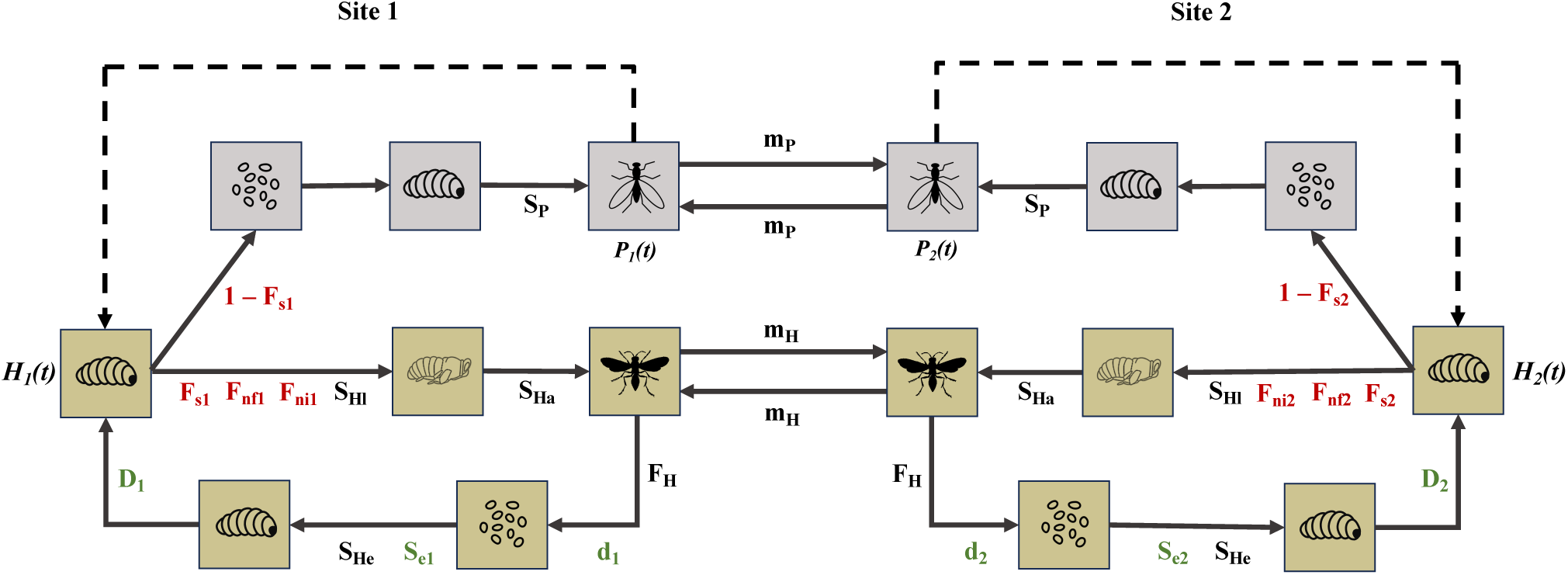
Schematic representation of the two-site dynamic of the *D. kuriphilus – T. sinensis* interaction integrating bottom-up, top-down regulations factors and dispersal. Parameters describing the basic life cycle of each species appear in black, while represented in red and green represent the effects of top-down control (hyperparasitism by native insects and fungi) and bottom-up control (chestnut tree frequency, density and genetic susceptibility), respectively. The parameters appearing in red and green are defined with respect to the site specific level of top-down and bottom-up control and therefore appear with index 1 or 2 specific to the modelled site.

To complete this original modelling, the definition of the quantities F_S_, F_ni_, F_nf_ and D were specified according to the available data on *D. kuriphilus* interactions with its hyper-parasites and knowledge about the competition for space between its larvae during galls formation. Assuming, as in the original Nicholson-Bailey model (64), a Poisson distribution of the parasitoid eggs in the host larvae, which account for the low ability of *T. sinensis* to fly towards galls of *D. kuriphilus*, the probability of escaping parasitism (F_s_) reads e^!“!#“$(&)^ where a_)_ stands for *T. sinensis* searching area. The probabilities of escaping native hyper-parasites (F_ni_ and F_nf_) were modelled as constant forcing terms as their population dynamics remain primarily determined by their native hosts. Finaly, the density-dependent survival of *D. kuriphilus* larvae (D) was modeled using a model proposed by Brännström and Sumpter (66) to account for such a typical ‘contest’ competition where winning individuals get all the (space) resources (required to make their chamber) while the other ones die. The survival D then reads K(1 − e^!*’(&)/-^) where H’(t) denotes the number of competiting *D. kuriphilus* larvae and K stands for the maximal population size, which corresponds to the product N.p_c_.k where N and k represent the density of trees in the forest site and the maximal amount of *D. kuriphilus* larvae that can be sustained per chestnut tree, respectively. The *C. sativa - D. kuriphilus - T. sinensis* interactions represented in Figure 1 then led to a non-spatial model defined as a pair of difference equations predicting the yearly changes in the abundance of *D. kuriphilus* larvae, H(t), and *T. sinensis* adults, P(t) :

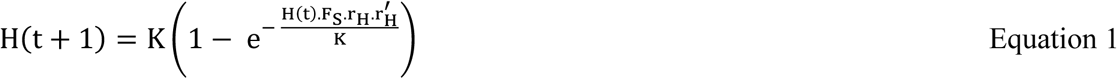

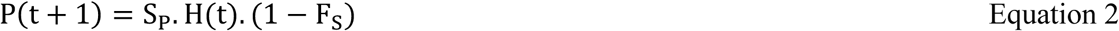

where r_*_= F_ni_.F_nf_.S_Hl_.S_Hp_ accounts for all survival rates of *D. kuriphilus* having escaped *T. sinensis* hyperparasitism until they become adults, and 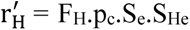 combines the adult fertility and the rates of eggs deposition and survival that determine the production of larvae entering into dormancy to overwinter (see Figure 1).

*The heterogeneity between sites and dispersal of the pest and its control agent.* The two sites model was designed to account for the between sites variations in parameters characterizing the bottom-up (i.e. N, p_c_, and S_e_) and top-down (i.e. F_ni_ and F_nf_) determinants of *D. kuriphilus* population dynamics, and for the dispersal of the pest and its control agent. The dispersal of these two species is restricted to the adult stage and their ability to navigate towards their hosts is limited to a small spatial scale (67,68). Accordingly, the two sites were coupled by a diffusive and symmetrical dispersal of *D. kuriphilus* and *T. sinensis* adults at rate m_H_ and m_P_, which led to two pairs of non-linear difference equations describing the annual changes in the population size of *D. kuriphilus* and *T. sinensis* in the first forest site;

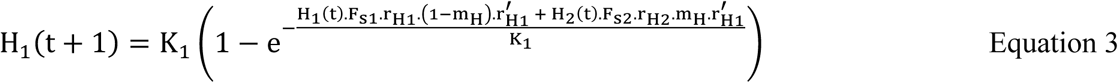

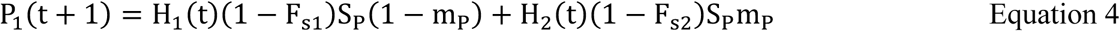

and in the second site,

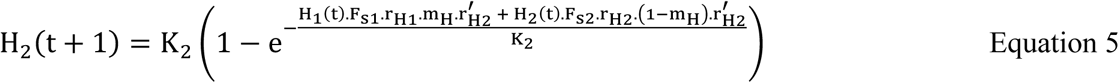

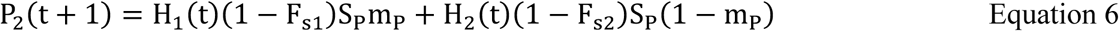

In equations 3-6, the quantities 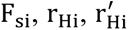 and K_i_varies between sites (i.e. with i=1,2) according to the specific values taken by the parameters that are involved in their formal definition (see above) and characterize the bottom-up (i.e. N, p_c_, and S_e_) and top-down (i.e. F_ni_ and F_nf_) determinants of *D. kuriphilus* local population dynamics.

### Analysis of the within site dynamic of tri-trophic interactions and site specific predictions

We first aimed at completing our understanding of the within site dynamic of *C. sativa* - *D. kuriphilus* - *T. sinensis* interactions by analysing the effect of the absolute density of forest trees (N), as it potentially is a further determinant of *D. kuriphilus* bottom-up control that showed a significant level of spatial heterogeneity between our 23 forest study sites located in the French Eastern Pyrenees. We expanded our previous analysis (24) by performing a local stability analysis of the within-site dynamics described by equations 1-2 (SM1). This analysis allowed to identify formal expressions of the conditions for i) *D. kuriphilus* to be able to invade, ii) *T. sinensis* to develop once its host has spread, and for iii) *D. kuriphilus* and *T. sinensis* to coexist at a stable equilibrium or through a regime of persisting oscillations. These conditions were used to represent the expected outcome of the tritrophic interaction in the frequency of chestnut trees (p_c_) - genetic susceptibility (Se) parameter plan drawn for three values of N corresponding to the minimal (N_min_=250), maximal (N_max_=1570) and average (N̅ =989) number of trees per hectare that we observed across our 23 forest sites. The conditions i) to iii) were then evaluated numerically by using the ‘nleqslv’ R package implemented in R.4.4.2 (69), and while all other parameters were set to their average value observed across the French Eastern Pyrenees (SM2).

We then aimed to use the within-site model to predict the intrinsic growth rates of *D. kuriphilus* (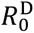) and *T. sinensis* (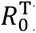) and the rate of *D. kuriphilus* biological control (C) in each of the 23 forest sites studied in the French Eastern Pyrenees, in order to compare them with the observed site specific rates of *D. kuriphilus* infestation and *T. sinensis* hyperparasitism. The intrinsic growth rate of *D. kuriphilus* was derived from equation 1 while considering no *T. sinensis* population, i.e. P(t)=0 so that F_s_=1, and the intrinsic growth rate of *T. sinensis* was obtained from equation 2 with the *D. kuriphilus* host population set at its equilibrium level (SM1). This led to the following expressions:

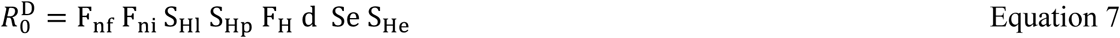

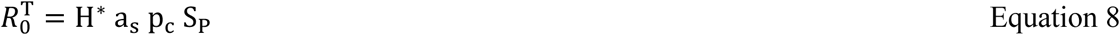

The rate of *D. kuriphilus* biological control C was defined as the percentage of reduction in the abundance of *D. kuriphilus* induced by *T. sinensis* and calculated from the dynamical equilibrium reached in the absence and presence of the control agent that were evaluated numerically from equations 1-2 using the ‘nleqslv’ R package implemented in R.4.4.2 (69). Quantitative estimates of those intrinsic growth (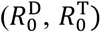) and control (C) rates were derived for each of the 23 forest sites by using the parameter values characterizing the bottom-up (i.e. N, p_c_, and S_e_) and top-down (i.e. F_s_, F_ni_ and F_nf_) determinants of *D. kuriphilus* population dynamics in each site. All other parameters were set to their average value observed across the French Eastern Pyrenees (SM2). Those predictions were then compared to the rates of *D. kuriphilus* and *T. sinensis* infestation observed across the 23 sampling sites (24). The correlation between the predicted intrinsic growth rates of *D. kuriphilus* (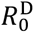) and the rates of chestnut tree infestation (measured in 2019 and 2020) was tested using a generalized linear mixed modelling (GLMM) that was conducted by following the procedure recommended by Zuur et al. (70). The impact of the sampling year (on the infestation measure considered) was characterized through a standard GLM analysis, and we subsequently accounted for such a temporal structure by incorporating sampling year as a random factor in our GLMM analyses (70, p.323-332). This statistical analysis was conducted in R.4.4.2 (69) using the glmmPQL function from the MASS package. The correlation between the predicted intrinsic growth rates of *T. sinensis* (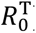) and the rates of hyperparasitism (observed in 2020), as well as the correlation between the predicted rates of biological control (C) and the reduction in the number of *D. kuriphilus* galls (observed in 2019 and 2020) were assessed using spearman’s rank correlation tests.

### Analysis of the two-sites dynamic of tri-trophic interactions and site specific predictions

The heterogeneity observed in the ‘bottom-up’ and ‘top-down’ control of *D. kuriphilus* population dynamics and in the above predictions of the within-site dynamic of interactions led to design the two-sites model corresponding to equations 3-6 to investigate the spatial dynamics that could result from such an environmental heterogeneity and the dispersal of *D. kuriphilus* and *T. sinensis* between sites. This model was analyzed by performing a local stability analysis of the two-sites dynamics described by equations 3-6 (SM3). The identified stability conditions were checked numerically using the ‘nleqslv’ R package (69). The characteristics of the forest environment (N, p_c_ and Se) in each site were then adjusted to their estimates derived from our 23 sampling sites, in order to investigate levels of heterogeneity corresponding to those observed in the French Eastern Pyrenees. The rate of *D. kuriphilus* and *T. sinensis* dispersal were then varied from 0 to 50% to test the effect of a large range of coupling between the various combinations of heterogeneity between the two sites.

The spatial expansion of the *C. sativa - D. kuriphilus - T. sinensis* model allowed to investigate the outcomes of their interactions, as well as the variations in control rate efficiencies at local and global scales, when pairs of sites from the same locality are connected through the dispersal of both the pest and its control agent. We coupled sites with similar (Figure 4,5A,B) or dissimilar (Figure 4,5C-H) local population dynamics predicted, using the combinations of forest density and frequency and genetic susceptibility of chestnut trees that we observed in the studied 23 forest sites. When coupling sites with stable and oscillatory co-existence dynamics, we performed a GLM statistical analysis to uncover the complex conditions for the two species to coexist at the two-site scale. Following the procedure recommended by Zuur et al. (70), we tested which of the bottom-up and top-down factors in both connected sites best explain the variations in the proportions of global stable and oscillatory co-existence dynamics.

**Figure 4.**
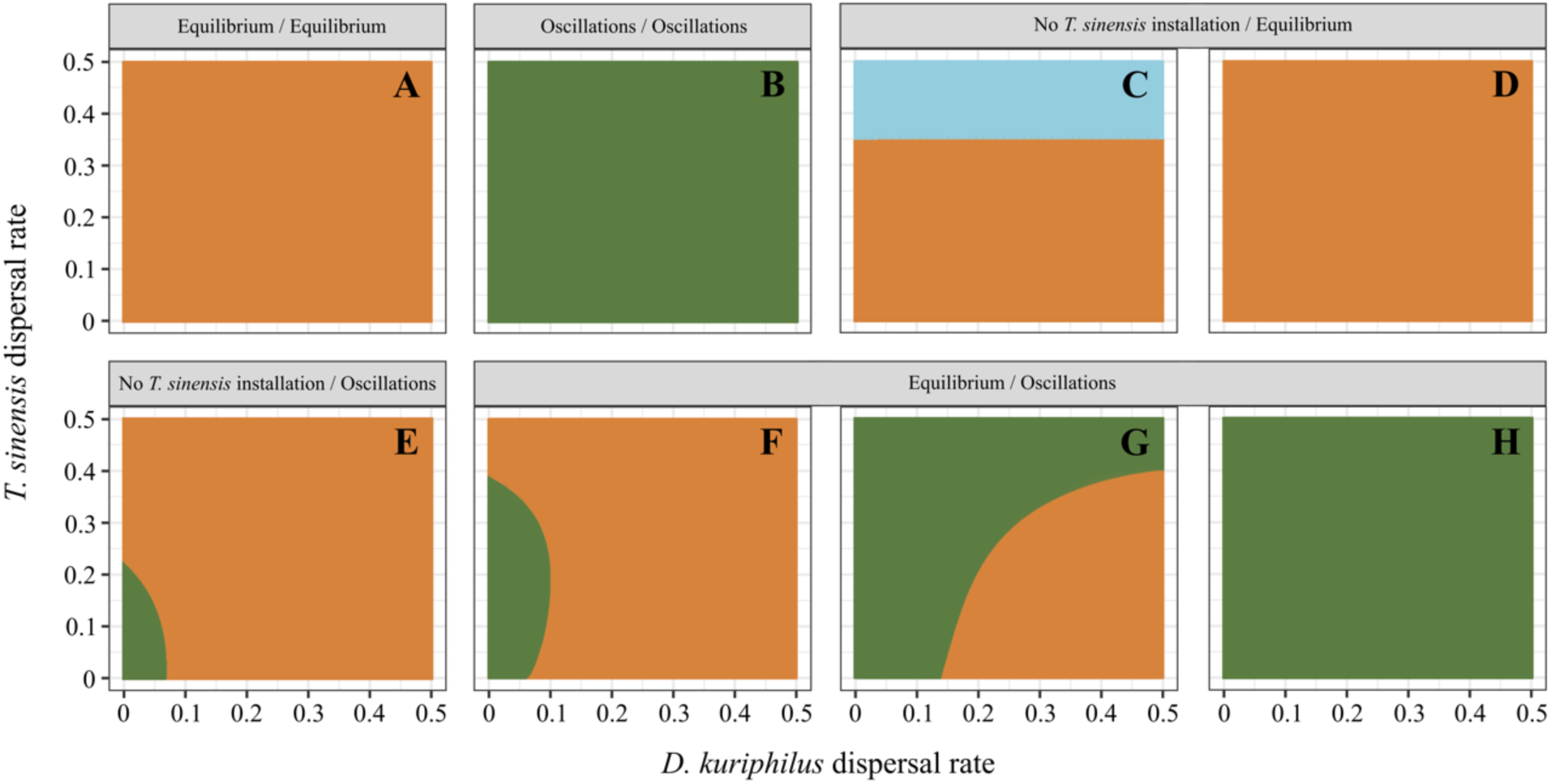
Dynamics of the *C. sativa - D. kuriphilus - T. sinensis* interaction in heterogeneous environments coupled *D. kuriphilus* and *T. sinensis* dispersal. The dynamic was predicted for combinations of sites with similar (A-B) or dissimilar (C-H) population dynamics predicted at the local scale, and while *D. kuriphilus* (x-axis) and *T. sinensis* (y-axis) rate of dispersal were varied from 0 to 50%. Colors indicate the nature of the predicted dynamic as in Figure 2A; coexistence of the two species in a stable (orange) or oscillatory (green) dynamics, *D. kuriphilus* invasion and failure of *T. sinensis* to persist (light blue).

**Figure 5.**
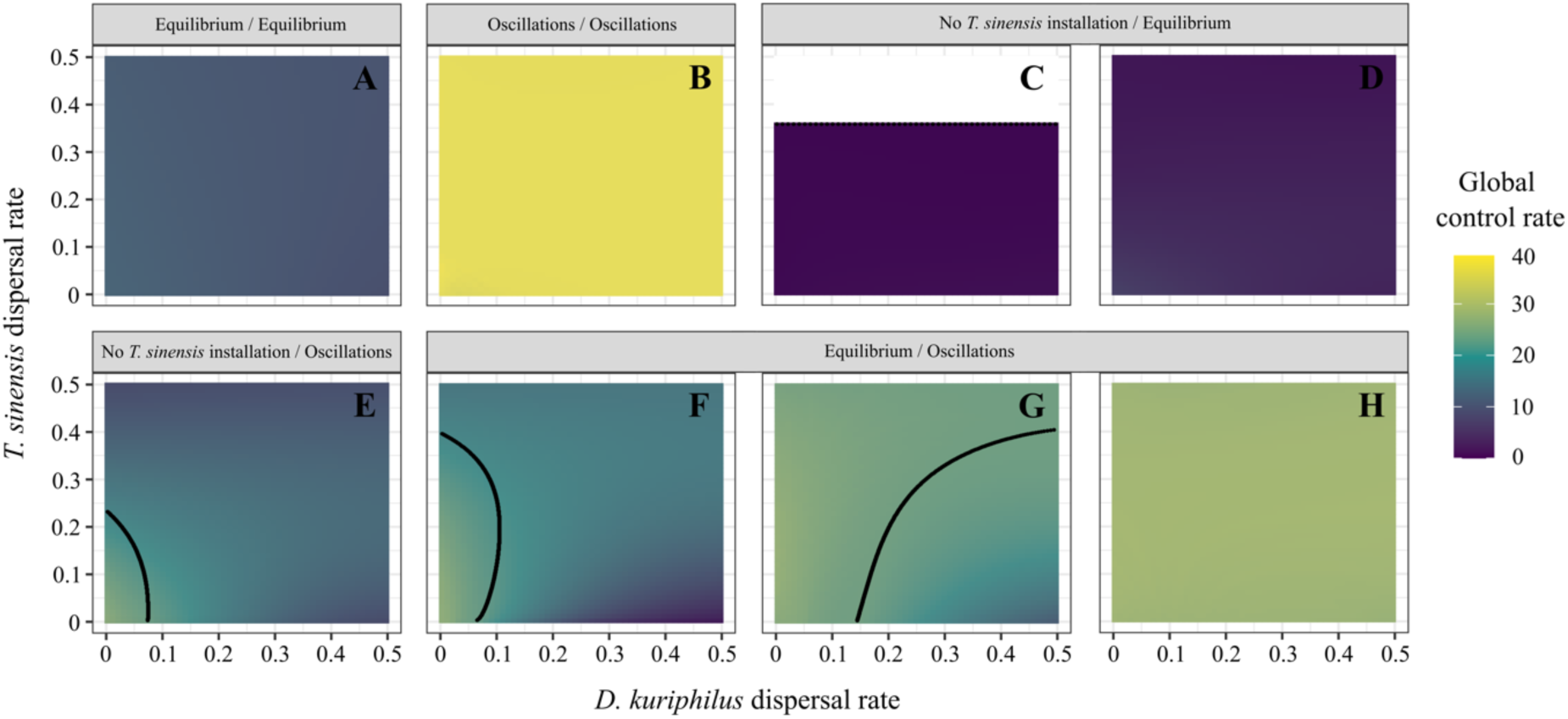
Variations in global biological control efficiency with respect to the dispersal of *D. kuriphilus* and *T. sinensis*. Global rate of biological control was predicted for the same combinations of sites as in Figure 4, with similar (A,B) or dissimilar (C-H) population dynamics predicted at the local scale, and while *D. kuriphilus* (x-axis) and *T. sinensis* (y-axis) rate of dispersal were varied from 0 to 50%. Colors indicate the variations in the predicted global control rate under the influence of *D. kuriphilus* and *T. sinensis* dispersal as represented in the legend. The black lines delineate the conditions where *D.kuriphilus* and *T. sinensis* co-exist with stable or oscilatory dynamics (E-G).

## Results

### Analysis of the within site dynamics of C. sativa - D. kuriphilus - T. sinensis interactions

The local stability analysis of the non-spatial model (SM1) unravelled 4 possible interaction dynamics according to the proportion of chestnut trees present in the forest stand (p_c_) and to their genetic susceptibility (Se). As expected, *D. kuriphilus* is unable to invade sites with low frequency and/or genetic susceptibility of chestnut trees (A-dark blue, B). When the frequency of chestnut trees increases, the pest starts spreading and reaches a low equilibrium level that does not allow for the control agent to establish itself (A-light blue, C). Above a threshold frequency of chestnut trees p_c_ of ∼40%, *D. kuriphilus* density is large enough for *T. sinensis* to spread, which lead to their coexistence in a stable way in mixed stands (A-orange, D) or through oscillatory dynamics in chestnut trees dominated forest sites, i.e. for p_c_ larger than ∼75% (A-green, E).

The overall density of trees had a significant impact on the outcome of the interaction between *D. kuriphilus* and *T. sinensis* when it was varied in the range of values observed across the 23 forest sites studied in the French Eastern Pyrenees (Figure 2, SM4). Increasing this density from its average to its maximal value lowered the threshold frequency of chestnut trees allowing for *T. sinensis* to persist from ∼40% to ∼30% (dotted lines) and the threshold above which the densities of *D. kuriphilus* and *T. sinensis* are expected to oscillate from ∼75% to ∼55% (continuous lines). Meanwhile, when the density of trees was lowered to its minimal value, the establishment of *T. sinensis* required a frequency of chestnut trees larger than ∼80% and this always led to a stable coexistence with *D. kuriphilus* as the abundance of chestnut trees could never be large enough for an oscillatory dynamic to develop, even if the corresponding site was to be a pure stand of *C. sativa*., i.e. p_c_ = 100%.

**Figure 2.**
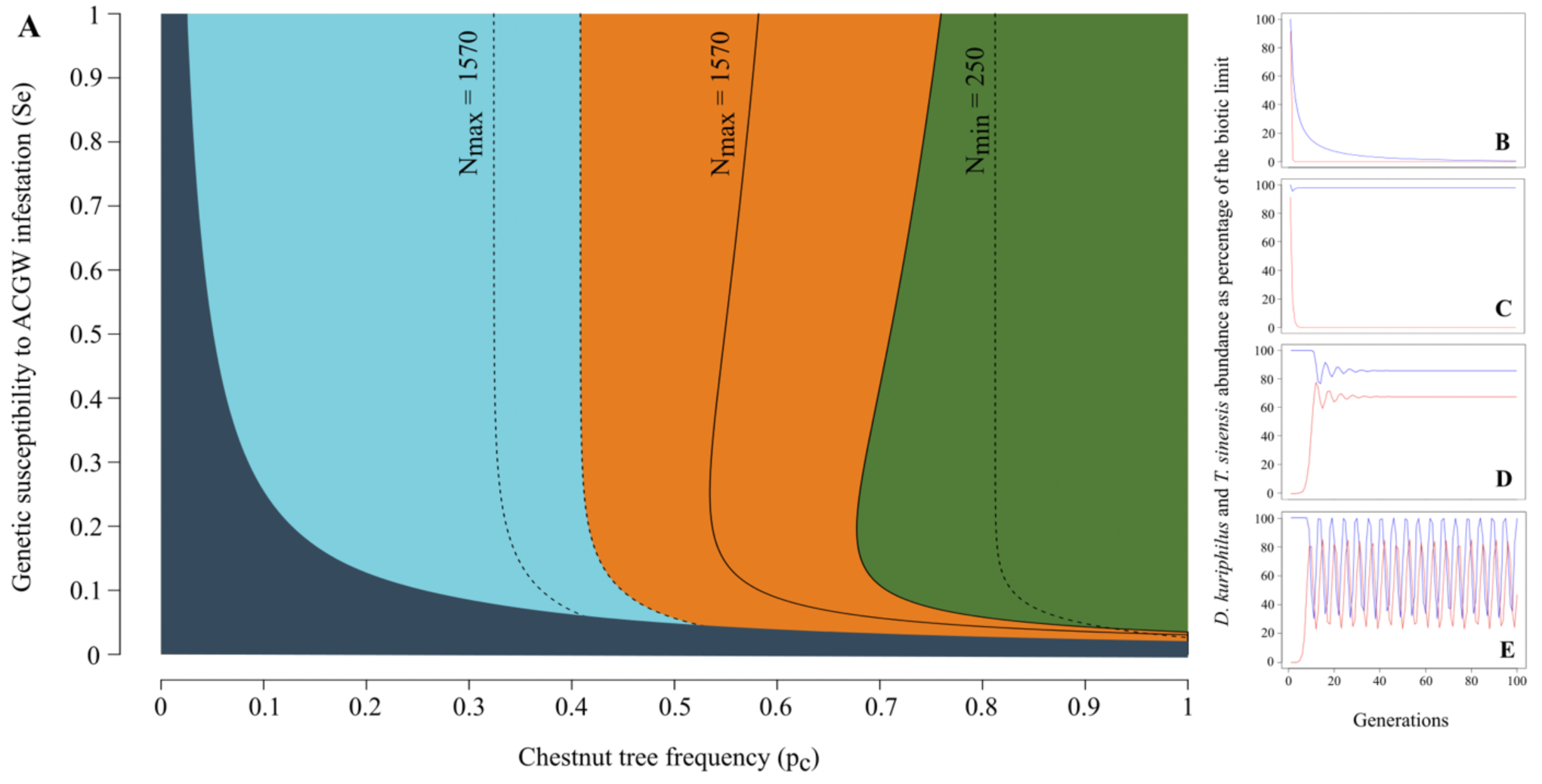
Effects of bottom-up determinants on the population dynamics of *D. kuriphilus - T. sinensis* interactions. (A) Dynamical outcomes of the population dynamic of *D. kuriphilus* and its control agent, *T. sinensis*, according to the frequency and genetic susceptibility of chestnut trees, and to the tree density in the forest site. Predicted dynamics: *D. kuriphilus* failure to invade (A-dark blue, B), *D. kuriphilus* invasion and failure of *T. sinensis* to spread (A-light blue, C), and coexistence of the two species in stable (A-orange, D) or oscillatory (A-green, E) dynamics. Dotted lines delineate the areas where *T. sinensis* is not able to persist (left) and where it spreads (right), while continuous lines indicate when *D. kuriphilus* and *T. sinensis* coexist in a stable way (left) and through oscillations (right). Those lines limit the blue, orange and green areas displayed for the average density of trees, i.e. 989 trees.h^-1^ (used in 24), and they are shifted to the left and to the right when the overall density of trees takes on the minimal (N_min_=250) and maximal (N_max_=1570) values that we observed across our 23 forest sites.

### The one-site model fails to explain the presence of T. sinensis in ‘sink’ sites with low abundance of chestnut trees

The one-site model was used to predict the intrinsic growth rate of *D. kuriphilus* (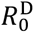) and *T. sinensis* (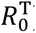) and the dynamics of their interaction with respect to the tree abundance, chestnut tree frequency and genetic susceptibility estimated in each of our 23 forest study sites (Figure 3A). Those predictions were then compared to the rates of chestnut tree infestation, i.e. the mean number of *D. kuriphilus* galls per bud, and to the rates of *D. kuriphilus* infestation, i.e. the proportion of *D. kuriphilus* larvae hyper-parasitized by *T. sinensis,* that we observed in the same field study sites. The observed rates of chestnut tree infestation were found significantly correlated with the *D. kuriphilus* growth rates (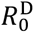) predicted from the bottom-up determinants measured in the forest sites (Figure 3B, GLMM analysis; χ^2^ = 0.06, df = 43, p = 0.0016). The observed rates of *D. kuriphilus* infestation by *T. sinensis* were also found significantly correlated with the predicted *T. sinensis* growth rates (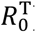) (Figure 3C, Spearman’s rank rho = 0.559, p = 0.016). Despite of these positive correlations, it is worth noting that i) *T. sinensis* was present at (lower but) significant rates in sites with a *T. sinensis* 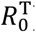 < 1 (Figure 3C), which resulted in a ii) lack of correlation between the observed reduction of *D. kuriphilus* between 2019 and 2020 (at a time where galls of the invasive were heavily infected by *T. sinensis,* Figure 3A) and the predicted rates of its control (Figure 3D, Spearman’s rank rho = 0.142, p = 0.517). We then hypothesized that the observation of *T. sinensis* in ‘sink’ sites, where it was predicted to be unable to establish because of the too low abundance of chestnut trees, could be explained by the dispersal of the control agent from the ‘source’ sites, where 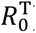 was found to reach a value of up to ∼3.

**Figure 3.**
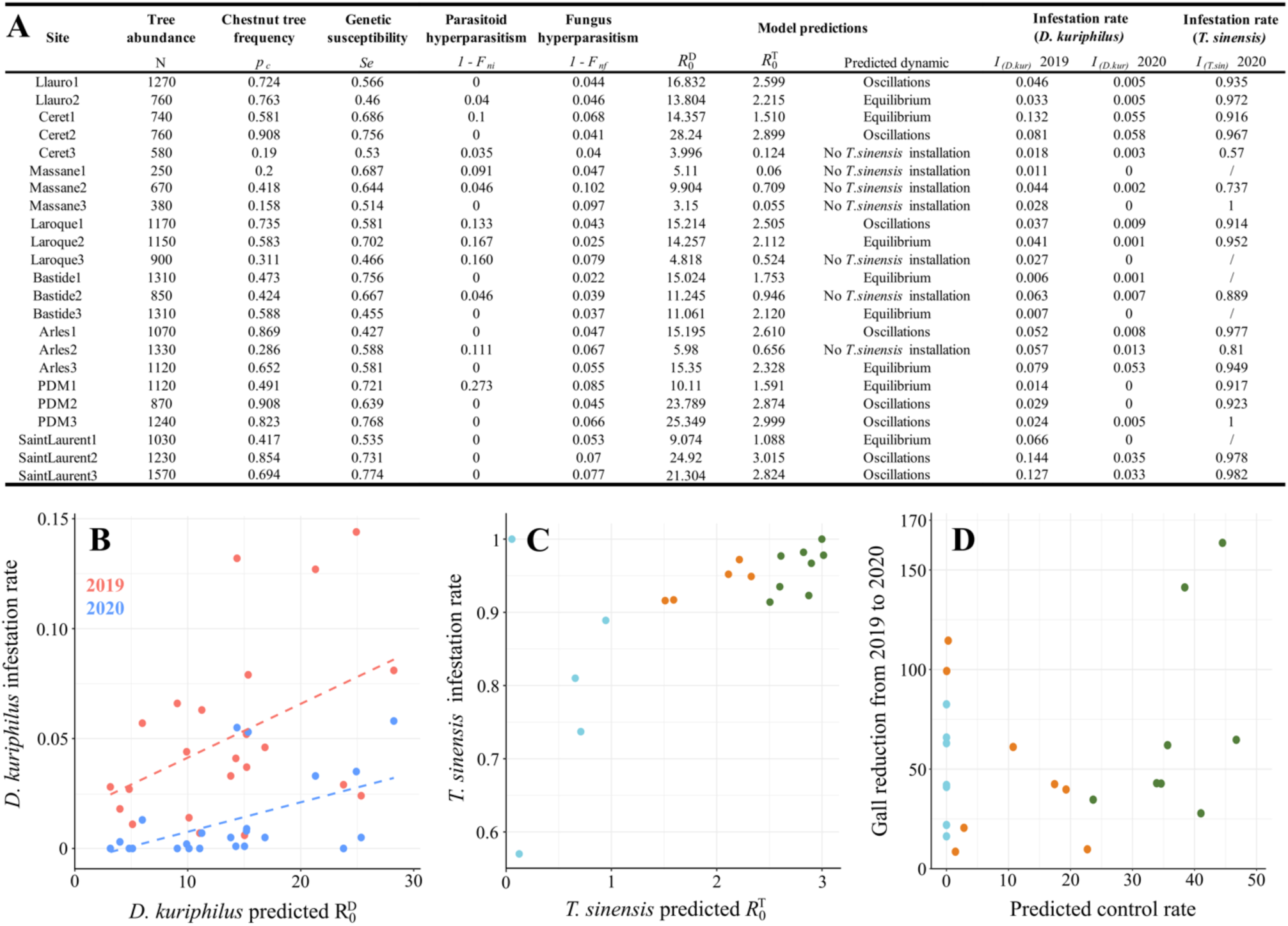
Predictions of the one-site model and comparisons with the rates of chestnut tree infestation by *D. kuriphilus* and *T. sinensis* hyperparasitism observed in the 23 forest sites. (A) Summary of the bottom-up and top-down determinants measured in each forest site, model predictions and observed rates of infestation by *D. kuriphilus* and their rates of hyperparasitism by *T. sinensis*. (B) Correlation between the predicted *D. kuriphilus* intrinsic growth rates (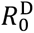) and the rates of chestnut tree infestation observed in 2019 (red) and 2020 (blue). (C) Correlation between the *T. sinensis* intrinsic growth rates (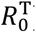) and the rates of hyperparasitism observed in 2020. (D) Correlation between the predicted rates of biological control and the gall reductions observed between 2019 and 2020. In C and D, each dot is colored according to the expected *D. kuriphilus* - *T. sinensis* population dynamics in the site, using the same color code as in Figure 2A.

### The dispersal of D. kuriphilus and T. sinensis synchronize their interaction dynamics in heterogeneous habitats

The spatial expansion of the *C. sativa - D. kuriphilus - T. sinensis* model allowed to investigate the outcomes of their interactions when two sites are connected through the dispersal of both the invasive pest (*D. kuriphilus)* and its control agent *(T. sinensis)*. We coupled sites with similar or dissimilar local population dynamics predicted by using the combinations of forest density and frequency and genetic susceptibility of chestnut trees that we observed in the 23 studied forest sites. The results obtained from all these combinations (SM5) led to 8 typical patterns for the possible outcomes of the *D. kuriphilus - T. sinensis* interaction with respect to their dispersal rates (Figure 4). The coupling of sites with the same local dynamics did not induce any qualitative changes at the local scale and, whatever the rates of dispersal of the invasive pest and its control agent, the global dynamics was therefore systematically similar to those predicted in each site (Figure 4A,B). More interestingly, when dispersal is introduced between sites exhibiting different local population dynamics, those dynamics tend to become synchronized and typically reach a stable (see orange parts of Figure 4C-G) or oscillatory (see green parts of Figure 4G,H) co-existence as soon as the parasite and control agent’s rates of dispersal are significantly increased.

Dispersal between a site where the two species coexist in a stable way and where the control agent cannot persist lead to the synchronization of the local dynamics of interaction with the stable persistence of the two species in both sites (Figure 4C,D). However, the highest rates of *T. sinensis* dispersal are detrimental to the control agent as its production in the source site where it can be sustained by a large enough population of *D. kuriphilus* is lost in the sink site where it cannot persist without dispersal.

The connection of a site where the two species coexist with an oscillatory dynamics and a site where i) the control agent cannot persist (Figure 4E) or ii) they coexist in a stable way (Figure 4F-H) can lead to similar changes in the outcome of the *D. kuriphilus* - *T. sinensis* interaction with respect to their dispersal rates. A rate of pest dispersal lower than 5-10% typically allows for the site exhibiting oscillations to destabilize the site with an intrinsically stable local dynamics while, at larger rates, the two local dynamics tend to converge towards the stable persistence of the two species in both sites (Figure 4E,F). The dispersal of the control agent (along the y-axis) induces similar dynamical changes, although larger rates of such a dispersal is required for the local dynamics to reach a stable equilibrium in each of the two sites (Figure 4E,F). However, coupling sites where the two species coexist in a stable and in an oscillatory way can also lead to a global oscillatory dynamics whatever their rates of dispersal (Figure 4H), or converge toward a stable equilibrium when *D. kuriphilus* dispersal rate is larger than 15-20% and *T. sinensis* dispersal rate remains lower than 30-40% (Figure 4G). According to our GLM analysis (SM6), the proportion of global oscillatory co-existence dynamics (Green parts in Figure 4F-H) increases with the tree community density and the chestnut tree frequency in both sites.

Overall, the dispersal of *D. kuriphilus* and *T. sinensis* between sites that differ in their forest structure (density, frequency and genetic susceptibility of chestnut trees), tend to synchronize the local dynamics of interaction. While those dynamics first coincide with the dynamics of the site with the highest chestnut tree frequency, they generally converge toward a stable coexistence of the pest and its control agent at larger rates of dispersal. Importantly, this led to the systematic persistence of *T. sinensis* in ‘sink’ sites when coupled with ‘source’ sites where the control agent is able to persist with its *D. kuriphilus* host either through stable (Figure 4C,D) or oscillatory (Figure 4E) dynamics.

### Dispersal between heterogeneous sites tends to homogenize the local efficiencies of biological control while decreasing its global rate

We lastly aimed to investigate how the changes in the *D. kuriphilus* - *T. sinensis* populations dynamics induced by their dispersal impact the local and global efficiencies of biological control of the invasive pest. Such rates of control were systematically evaluated for all combinations of sites (SM7) and the variations of the global rate averaged over the two sites are shown in Figure 5 for the same representative pairs of sites as in Figure 4. The dispersal between sites with similar local dynamics did not induce any significant changes in the biological control rates at both local (SM8) and global (Figure 5A,B) scales, whatever the rates of dispersal of *D. kuriphilus* and *T. sinensis*. The global efficacy was 3-4 times larger when coupling sites where the invasive pest and its control agent coexist through an oscillatory dynamics than where they persist in a fixed equilibrium. Such global control rates indeed typically reached ∼40% and ∼13%, respectively. The coupling of sites exhibiting dissimilar local dynamics (Figures C-H) tend to reduces the differences in local efficiencies of biological control (SM8). As expected, when *T. sinensis* is not expected to persist in one of the sites (Figure 5C-E), the global control typically took on low values. At local scales, the coupling between such *T. sinensis* sink and source sites indeed allows for the installation of *T. sinensis* in the former at the expense of the local control in the latter (SM8). Such a balance can actually lead to the complete lost of control at high rate of *T. sinensis* dispersal (Figure 5C). Meanwhile, the coupling with a site exhibiting *D. kuriphilus - T. sinensis* oscillations can also truly allow for *T. sinensis* to spread and persist into a sink environment, albeit at low abundances (Figure 5E). Finally, the global efficacy of control take on intermediate values when the pest and its control agent are expected to persist in both sites through a fixed equilibrium and oscillations (Figure 5F-H). Such global rate of control ranged between 5% and 35% depending on disperal rates, and the local efficiency of control then converge to similar values as dispersal rates increase (SM8). Overall, across all those predicted patterns (Figure 5A-H), the dispersal of *D. kuriphilus* consistently reduced the global rate of biological control, while *T. sinensis* dispersal effect could vary from detrimental to beneficial when *D. kuriphilus* dispersal increases (Figure 5E-G).

## Discussion

A cornerstone of research about plant pest biological invasion remains the difficult untangling of the bottom-up and top-down forces that shape their dynamics (e.g. 71,72). While such an understanding is critical to identify the key determinants of pest invasion and choose adequate control strategies (73,74), most studies usually do not account for the spatial heterogeneity in such forces and the dispersal between sites that can represent ‘source’ or ‘sink’ for the invasive pest and/or its biological control agent (75, and references therein). By combining the predictions of an integrative within-site model of *D. kuriphilus* invasion and its bottom-up and top-down regulation forces with those of its extension into a two-site model, we investigated the ‘source-sink’ dynamics induced by the dispersal of this invasive pest and its control agent, *T. sinensis*, in spatially heterogeneous forest environments encountered in the French Eastern Pyrenees (24).

The well-parameterized within site modelling we designed to integrate the outcomes of previous field and genomic studies (24), provided an unprecedented set of predictions about the dynamics of *D. kuriphilus* - *T. sinensis* interaction according to the observed characteristics of the forest environment, i.e. the tree density and chestnut tree frequency and the genetic susceptibility to *D. kuriphilus* infestation, in each of the 23 sites sampled in the French Eastern Pyrenees. The composition of the forest tree community showed strong variations among those sites that were significantly associated with heterogeneous levels of *D. kuriphilus* infestation in 2019 (χ^2^ = 479.8, df = 22, p < 2.2 10^-16^) and 2020 (χ^2^ = 428.1, df = 22, p < 2.2 10^-16^). The model predicted that the forest tree structure should lead to the coexistence of *D. kuriphilus* and *T. sinensis* in 16/23 sites, where both species were indeed observed in both 2019 and 2020. Interestingly, the prediction of the persistence of a control agent and its target does not fit the classical theory of biological control whereby the introduced agent leads to the extinction of the pest followed by its own (e.g. 76). It is however consistent with i) previous outcomes of the modelling of this biological system by Paparella et al. (77) and Zitoun et al. (24) that showed the *D. kuriphilus* - *T. sinensis* tendency to coexist by exhibiting stable equilibrium or oscillations in abundances, and with ii) field observations that the introduction of *T. sinensis* did not allow for the elimination of the pest (55) and instead led to three successive peaks of *D. kuriphilus* shortly followed by peaks of *T. sinensis* over 25 years of field interaction (57). Meanwhile, in the absence of dispersal, our modelling predicted that *T. sinensis* should not be able to spread in the remaining 7/23 ‘sink’ sites, where the control agent was nonetheless systematically observed. Such a discrepancy between the predictions of our integrative within site model and the field assessment of *T. sinensis* population pointed toward hidden source-sink dynamics (60) within the study area, and prompted the extension of our tritrophic modelling into a two-site model.

The spatial modelling provided the first predictions about the global dynamics resulting from the dispersal of both *D. kuriphilus* and *T. sinensis* between sites with different level of bottom-up and top-down controls, which allowed to assess the hypothesis that the presence of the control agent in ‘sink’ sampling site could be explained by its dispersal from surrounding ‘source’ sites. The primary outcome of the spatial model analysis is that, even at low rates of dispersal, the dynamics of the site with the lowest frequency of chestnut trees tend to converge to the dynamics of the site where the frequency is higher, which is consistent with its strong positive effect on *D. kuriphilus* abundance that we both predicted from the within-site model and observed in the field (24 and this study). Such an homogenization effect is also consistent with a large set of theoretical studies that have previously demonstrated the strong ability of dispersal to synchronize various ecological dynamics (78–80). Interestingly, when chestnut trees are the dominant species in the ’driving’ site, as observed in the ‘Ceret2’ and ‘PDM2’sites, the local dynamics in the other site and the global dynamics all exhibit oscillations as predicted in the ‘driving’ site, as long as *D. kuriphilus* dispersal rate remains low enough. Meanwhile, when the pest dispersal rate is increased, the local and global dynamics tend to converge toward a stable co-existence. While the dispersal rate of this species remains loosely quantified, they are usually thought to be low due to the short adult lifespan (62). Accordingly, one should expect large patches of host resources to sustain some oscillatory dynamics rather than the local and global dynamics to converge to a stable spatially homogeneous equilibrium. This seems to be supported by the recent observation of an re-increase in infestation in 17/23 of our sampling sites (unpublished data). A secondary key predictions of the spatial modelling is that *T. sinensis* population in ‘sink’ sites can systematically be sustained by dispersal from ‘source’ sites, whatever be the precise dynamical regime of *D. kuriphilus - T. sinensis* coexistence in the source. This provides a strong support to the dispersal hypothesis to explain the observation of *T. sinensis* in the 7 predicted ‘sink’ sites as the other (two) sampling sites located within the same stations, i.e. Arles sur Tech, Céret, Bastide and Laroque, were typically shown to be ‘source’ sites of *T. sinensis* with R_0_ values larger than 2 (see Figure 3). The only notable exception correspond to the ‘La Massane’ natural reserve, where all three sites were predicted to be ‘sink’. The spread of *T. sinensis* in this area with limited amount of chestnut trees can potentially be explained by dispersal from the neighboring station ‘Laroque’, where 2 of the 3 sampling sites were predicted to be sources of *T. sinensis* with large R_0_ values.

The two sites integrative model further allowed to investigate the effect of dispersal on the biological control efficiency while accounting for the observed level of heterogeneity in the forest environments encountered in the French Eastern Pyrenees. The effect of *T. sinensis* dispersal was found to be mostly detrimental to the overall rate of control that can be achieved across (the two) sites. Although *T. sinensis* dispersal from ‘sources’ allows for its persistence and therefore a biological control in ‘sink’ sites, this gain of control did not typically compensate for the decline of control observed in the source site due to *T. sinensis* emigration. The low capacity of the control agent to find its hosts in sites with unfavourable conditions indeed lead to a typical dilution effect (81) and therefore to a decline of the *T. sinensis* abundance across sites. This reduction of the global control efficacy was found to be particularly important for sites that exhibits large differences in chestnut tree abundance and therefore in their local control rates. On the contrary, an increase of the biological control efficiency with *T. sinensis* dispersal was shown to be possible when coupling two sites allowing for *D. kuriphilus* - *T. sinensis* coexistence, although such a gain required the local dynamic to exhibits oscillatory dynamics in at least one site with a high level of biological control. The dispersal between sites in oscillations dynamics then helped to a quicker reconstruction of the parasitoid populations, as suggested by Mohd and Noorani (82). Meanwhile, the magnitude of this effect always remained low, i.e. only rising up to +0.4%. Overall, *D. kuriphilus* dispersal was therefore shown to lead to a global control reduction that was stronger for larger chestnut tree abundance differences between connected sites.

The above results contrast with the predictions derived from the ‘Enemies Hypothesis’ theorized by Root (83), whereby spatial heterogeneity has a beneficial effect on the biocontrol of phytophagous pests. The diversification of plant species in the environment is then thought to allow for a better biocontrol via an increase in the number of prey or alternative hosts and in the proportion of over- wintering sites or refuges against disturbances for control agents (84,85). While this concept originally developed for agroecosystems has been transposed to the forest environment (86,87), it has also been pointed out that it is largely based on the assumption that predators are generalists (88). On contrary, specialist control agents, particularly parasitoids, tend to be more abundant and effective in homogeneous systems than in heterogeneous systems (89) as they depend on a single resource whose availability is impacted by the loss of favorable habitat (90), which explains that we predicted the global *T. sinensis* efficacy to decrease when chestnut tree forest sites are connected by dispersal.

## ACKNOWLEDGMENTS

This study is set within the framework of the ‘Laboratoires d’Excellences (LABEX)’ TULIP (ANR- 10-LABX-41) and of the ‘École Universitaire de Recherche (EUR)’ TULIP-GS (ANR-18-EURE- 0019). This work has benefited from a PhD fellowship to JLZ (Région Occitanie,) and from the project ‘Modélisation hybride d’invasions biologiques’ (MITI CNRS, PI. Gourbiere S.). This work further benefited from the modelling discussions hold in the context of the ‘PHYTOMICS’ (ANR, ANR-21- CE02-0026, PI. Gourbiere S.) and ‘ELVIRA’ (ANR, ANR-21-CE20-0041, PI. Piganeau G.) project. The funders played no role in the study design, data collection and analysis, decision to publish, or preparation of the manuscript.

