## Supplementary Materials for "Source-sink dynamics explains the co-existence of the invasive pest *Dryocosmus kuriphilus* and its biological control agent *Torymus sinensis* across French Eastern Pyrenees"

### SUPPORTING INFORMATION

**Supplementary Material 1.** Local stability analysis of the one-site *D. kuriphilus* – *T. sinensis* interaction model.

Consider the equations of the non-spatialized model:

$$\begin{cases} H(t+1) = K\left(1 - e^{-\frac{RH(t)e^{-\alpha'_s P_S(t)}}{K}}\right) \\ P_s(t+1) = cH(t)\left(1 - e^{-\alpha'_s P_S(t)}\right) \end{cases}$$

where $R = \mathrm{F_{nf}\,F_{ni}\,S_{Hl}\,S_{Hp}\,F_H\,d\ \ Se\,S_{He}}$ and c = $S_P$

The Jacobian matrix at equilibrium $H^*$ and $P_s{}^*$, is :

$$A_{(H^*,P^*)} = \begin{pmatrix} \frac{\partial f}{\partial H^*} & \frac{\partial f}{\partial P^*} \\ \frac{\partial g}{\partial H^*} & \frac{\partial g}{\partial P^*} \end{pmatrix} = \begin{pmatrix} Re^{-\frac{RH^*}{K}e^{-\alpha'_s P_S{}^*} - \alpha'_s P_S{}^*} & -RH^*\alpha'_s e^{-\frac{RH^*}{K}e^{-\alpha'_s P_S{}^*} - \alpha'_s P_S{}^*} \\ c(1 - e^{-\alpha'_s P_S{}^*}) & cH^*\alpha'_s e^{-\alpha'_s P_S{}^*} \end{pmatrix}$$

where $H^* = K\left(1 - e^{-\frac{RH^*}{K}e^{-\alpha'_s P_S{}^*}}\right)$, so that $e^{-\frac{RH^*}{K}e^{-\alpha'_s P_S{}^*}} = 1 - \frac{H^*}{K}$ leading to :

$$A_{(H^*,P^*)} = \begin{pmatrix} R(1 - \frac{H^*}{K})e^{-\alpha'_s P_S{}^*} & -RH^*\alpha'_s(1 - \frac{H^*}{K})e^{-\alpha'_s P_S{}^*} \\ c(1 - e^{-\alpha'_s P_S{}^*}) & cH^*\alpha'_s e^{-\alpha'_s P_S{}^*} \end{pmatrix}$$

From this matrix, the stability of any equilibrium points can be found using the stability condition that is equivalent to the absolute value of both eigenvalues of A being lower than 1 (91, page 57) :
$|tr(A)| < \det(A) + 1 < 2$.

1) We first considered the stability of the trivial equilibrium ($H^* = 0, P_s{}^* = 0$).

The stability condition then simplifies to R < 1 as tr(A)=R and det(A)=0. This leads to

$$\mathrm{F_{nf}\,F_{ni}\,S_{Hl}\,S_{Hp}\,F_H\,d\ \ Se\,S_{He}} < 1$$

which we expressed as

$$Se < \frac{1}{\mathrm{F_{nf}\,F_{ni}\,S_{Hl}\,S_{Hp}\,F_{H}\,d\;\;S_{He}}}$$

to ease its representation in Figure 1, where it defines the upper limit of the dark blue area.

2) We second considered the stability of the equilibrium where only *D. kuriphilus* persists, i.e. ($H^*$,0).

The Jacobian matrix then becomes

$$\mathrm{A}_{(H^*,0)} = \begin{pmatrix} R\left(1-\frac{H^*}{K}\right) & -RH^*\alpha'_s(1-\frac{H^*}{K}) \\ 0 & cH^*\alpha'_s \end{pmatrix}$$

and the stability condition reads:

$$\left|R\left(1-\frac{H^*}{K}\right)+\;cH^*\alpha'_s\right| < R\left(1-\frac{H^*}{K}\right)\;cH^*\alpha'_s + 1 < 2$$

In the absence of the control agent *T. sinensis*, i.e. $P_s{}^* = 0$, *D. kuriphilus* tends to saturate the environment : $H^* = K\left(1-e^{-\frac{RH^*}{K}}\right) \longrightarrow K(1-e^{-R}) \approx K$ (as R is typically in the range of 10-20, e.g. 24), and the condition simplifies as

$$cH^*\alpha'_s < 1$$

which can be re-written

$$cK(1-e^{-R})\alpha'_s < 1$$

and, since $R = \mathrm{F_{nf}\,F_{ni}\,S_{Hl}\,S_{Hp}\,F_{H}\,d\;\;Se\;S_{He}}$ , this leads to

$$Se < \frac{ln\left(1-\frac{1}{cK\alpha'_s}\right)^{-1}}{\mathrm{F_{nf}\,F_{ni}\,S_{Hl}\,S_{Hp}\,F_{H}\,d\;\;S_{He}}}$$

which allowed to draw the upper limit of the light blue area in Figure 1.

The last two dynamics to discriminate were when both species would coexist at a stable equilibrium ($H^*$,$P^*$) and when such point would be unstable and the two species would instead coexist through stable oscillations. The stability condition of the equilibrium point ($H^*$,$P^*$) read:

$$|R\,A\,B + cH^{*}\alpha'_{s}B| < \; R\,A\,B\,cH^{*}\alpha'_{s} + 1 < 2$$

where $A = R\left(1 - \frac{H^{*}}{K}\right)$ and $B = \; cH^{*}\alpha'_{s}$.
Those conditions were evaluated numerically, which allowed to draw the line between the orange and green areas in Figure 1.

**Supplementary Material 2**. Definition and estimates of the parameters of the *D. kuriphilus* host - *T. sinensis* and native parasitoids dynamical model.

| | Symbol | Estimate and 95% confidence interval | References |
|---|---|---|---|
| ***Dryocosmus kuriphilus*** | | | |
| Egg survival to hypersensitive response | $S_e$ | 0.680 | 24 |
| Egg survival | $S_{He}$ | 0.535 ± 0.003 | 46 |
| Larvae survival | $S_{Hl}$ | 0.936 ± 0.010 | 50 |
| Pupae survival | $S_{Hp}$ | 0.952 ± 0.010 | 48 |
| Fertility | $F_H$ | 90 ± 6 | 24 |
| Maximum carrying capacity per hectare | K | 14 880 000 | 24 |
| ***Torymus sinensis*** | | | |
| Larvae and adult survival | $S_P$ | 0.913 ± 0.004 | 92 |
| Searching area | $a_s$ | $4.4 \times 10^{-4} \pm 0.4 \times 10^{-4}$ | 24 |
| ***Native hymenoptera and fungi*** | | | |
| Probability to escape hymenopteran infection | $F_{ni}$ | 0.955 ± 0.02 | 24 |
| Probability to escape fungal infection | $F_{nf}$ | 0.946 ± 0.001 | 24 |

**Supplementary Material 3.** Local stability analysis of the two-sites *D. kuriphilus – T. sinensis* interaction model.

Considering the equations of the two-sites model;

$$\begin{cases} H_1(t+1) = K_1\left(1 - e^{-\frac{H_1(t).F_{s1}.r_{H1}.(1-m_H).r'_{H1} + H_2(t).F_{s2}.r_{H2}.m_H.r'_{H1}}{K_1}}\right) \\ P_1(t+1) = H_1(t)(1-F_{s1})S_P(1-m_P) + H_2(t)(1-F_{s2})S_P m_P \\ H_2(t+1) = K_2\left(1 - e^{-\frac{H_1(t).F_{s1}.r_{H1}.m_H.r'_{H2} + H_2(t).F_{s2}.r_{H2}.(1-m_H).r'_{H2}}{K_2}}\right) \\ P_2(t+1) = H_1(t)(1-F_{s1})S_P m_P + H_2(t)(1-F_{s2})S_P(1-m_P) \end{cases}$$

where $r_{Hi} = F_{nii}.F_{nfi}.S_{Hl}.S_{Hp}$ and $r'_{Hi} = F_H.p_{ci}.S_{ei}.S_{He}$,

the Jacobian matrix at the equilibrium $H_1^*$, $P_1^*$, $H_2^*$ and $P_2^*$, reads

$$A_{(H_1^*,P_1^*,H_2^*,P_2^*)} = \begin{pmatrix} A_{1.1} & A_{1.2} \\ A_{2.1} & A_{2.2} \end{pmatrix}$$

with :

$$A_{1.1} = \begin{pmatrix} R_{1.1}F_{s1}e^{-\frac{R_{1.1}H_1^*F_{s1}+R_{2.1}H_2^*F_{s2}}{K_1}} & -R_{1.1}H_1^*\alpha'_1F_{s1}e^{-\frac{R_{1.1}H_1^*F_{s1}+R_{2.1}H_2^*F_{s2}}{K_1}} \\ S_P(1-F_{s1})(1-m_P) & S_PH_1^*\alpha'_1F_{s1}(1-m_P) \end{pmatrix}$$

$$A_{1.2} = \begin{pmatrix} R_{2.1}F_{s2}e^{-\frac{R_{1.1}H_1^*F_{s1}+R_{2.1}H_2^*F_{s2}}{K_1}} & -R_{2.1}H_2^*\alpha'_2F_{s2}e^{-\frac{R_{1.1}H_1^*F_{s1}+R_{2.1}H_2^*F_{s2}}{K_1}} \\ S_P(1-F_{s2})m_P & S_PH_2^*\alpha'_2F_{s2}m_P \end{pmatrix}$$

$$A_{2.1} = \begin{pmatrix} R_{1.2}F_{s1}e^{-\frac{R_{1.2}H_1^*F_{s1}+R_{2.2}H_2^*F_{s2}}{K_2}} & -R_{1.2}H_1^*\alpha'_1F_{s1}e^{-\frac{R_{1.2}H_1^*F_{s1}+R_{2.2}H_2^*F_{s2}}{K_2}} \\ S_P(1-F_{s1})m_P & S_PH_1^*\alpha'_1F_{s1}m_P \end{pmatrix}$$

$$A_{2.2} = \begin{pmatrix} R_{2.2}F_{s2}e^{-\frac{R_{1.2}H_1^*F_{s1}+R_{2.2}H_2^*F_{s2}}{K_2}} & -R_{2.2}H_2^*\alpha'_2F_{s2}e^{-\frac{R_{1.2}H_1^*F_{s1}+R_{2.2}H_2^*F_{s2}}{K_2}} \\ S_P(1-F_{s2})(1-m_P) & S_PH_2^*\alpha'_2F_{s2}(1-m_P) \end{pmatrix}$$

where $R_{1.1} = r_{H1}(1-m_H)r'_{H1}$ and $R_{2.2} = r_{H2}(1-m_H)r'_{H2}$;

$R_{1.2} = r_{H1}m_Hr'_{H2}$ and $R_{2.1} = r_{H2}m_Hr'_{H1}$;

$F_{s1} = e^{-\alpha'_1P_1^*}$ and $F_{s2} = e^{-\alpha'_2P_2^*}$.

The submatrices of the Jacobian diagonal bring together the intra-site contributions ($A_{1.1}$ for site 1 and $A_{2.2}$ for site 2), while the remaining submatrices brings together the inter-site contributions. Therefore, in the absence of migration between sites (i.e.$m_H = 0$ and $m_P = 0$) $A_{1.1}$ and $A_{2.2}$ become similar to those obtained in the one-site model local stability analysis (see SM1), while $A_{1.2}$ and $A_{2.1}$ become zero matrix.

From the above Jacobian matrix, the stability of any equilibrium points can be found by demonstrating that all real eigenvalues of A are between -1 and 1 and that for all its complex eigenvalues ($\lambda = A + B_i$): $\sqrt{A^2 + B^2} < 1$ (93, page 319).

**Supplementary Material 4**. Predicted dynamics of the *D. kuriphilus – T. sinensis* interaction in each of the 23 studied sites according to the chestnut tree density, frequency and genetic susceptibility. For each of the 23 panels, the dot indicates the frequency (x-axis) and average genetic susceptibility (y-axis) of chestnut trees observed in the corresponding sampling site, while the limits between the different areas are set with respect to the observed tree density.

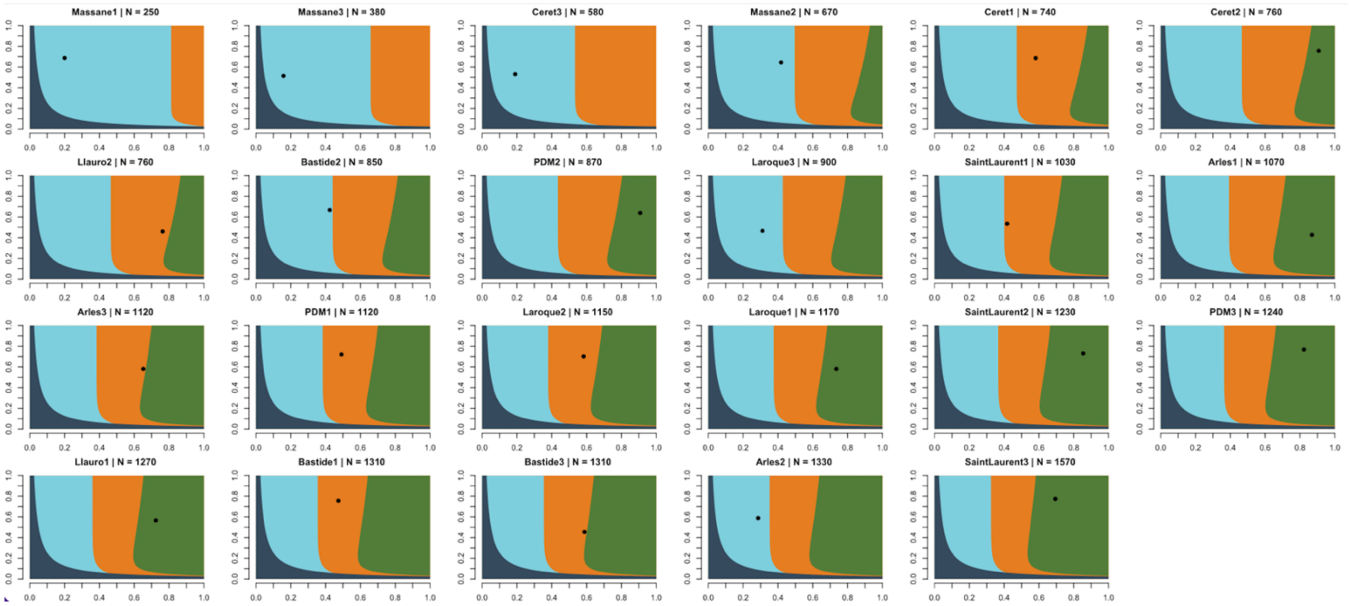

**Supplementary Material 5. Dynamics of the *D. kuriphilus* - *T. sinensis* interaction in heterogeneous environments made of two sites coupled by the dispersal of both species**. Dynamics were predicted for all combinations of sites within a locality, with similar (A-C) or dissimilar (D-S) population dynamics predicted at the local scale, and while *D. kuriphilus* and *T. sinensis* rates of dispersal were varied from 0 to 50%. Colors indicate the nature of the predicted dynamic as in Figure 2A; coexistence of the two species in a stable (orange) or oscillatory (green) dynamics, *D. kuriphilus* invasion and failure of *T. sinensis* to persist (light blue).

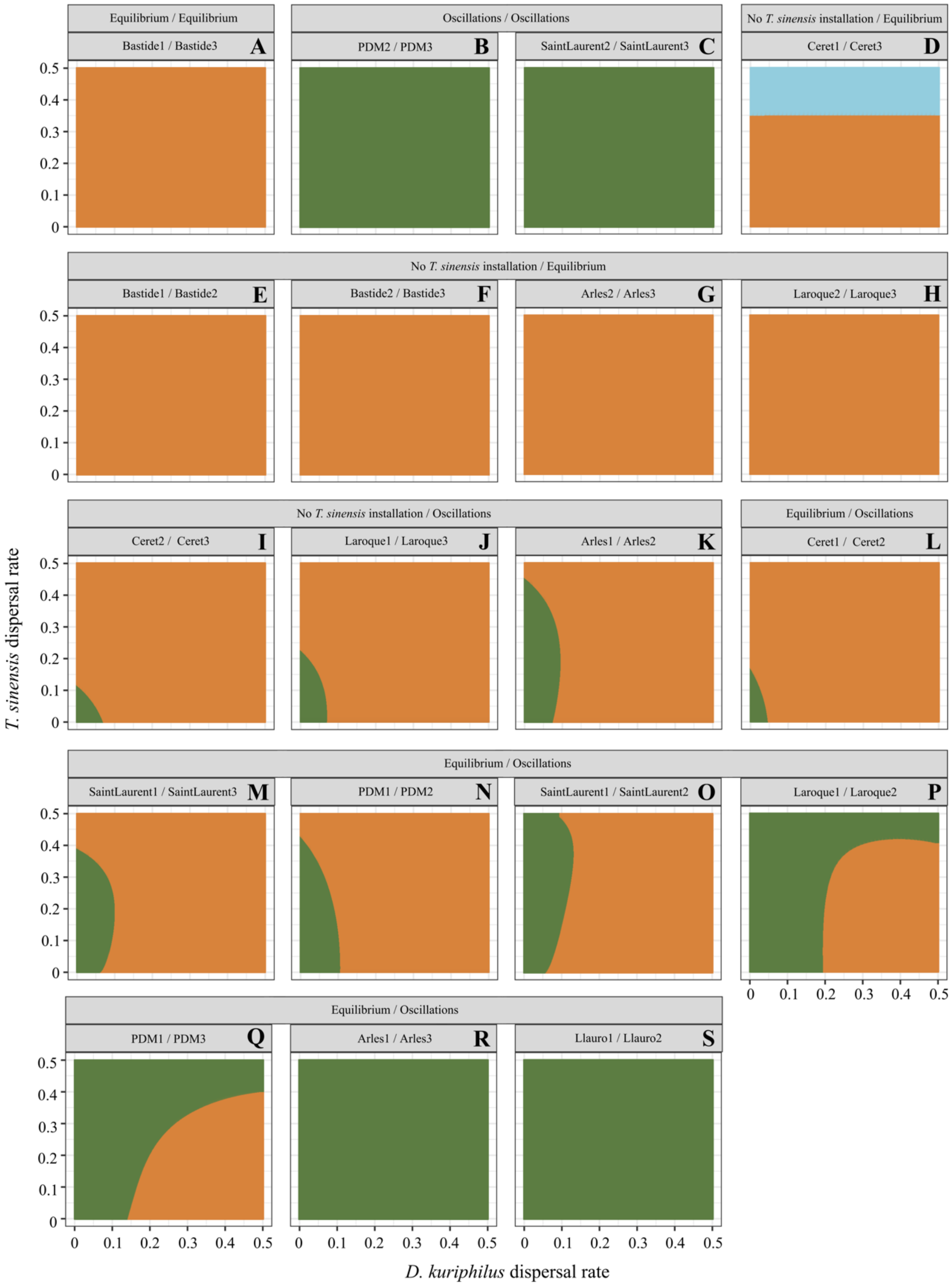

**Supplementary Material 6.** GLM analysis uncovering which of the bottom-up and top-down factors of each of the two connected sites best explain the variations in the proportion of oscillatory dynamics in the patterns predicted when coupling sites where *D. kuriphilus* and *T. sinensis* coexist through an oscillatory dynamics and in a stable way.

| Fixed effects | Estimate | Standard error | *df* | P |
|---|---|---|---|---|
| Response variable : Oscillatory dynamics proportions with dispersal | | | | |
| Intercept | -65.788695 | 10.203697 | 7 | 0.00757 |
| Tree density in stable site | 0.006527 | 0.001834 | 7 | 0.03784 |
| Chestnut tree frequency in stable site | 32.564671 | 4.609445 | 7 | 0.00583 |
| Tree density in oscillatory site | 0.014025 | 0.002382 | 7 | 0.00977 |
| Chestnut tree frequency in oscillatory site | 31.218387 | 6.165474 | 7 | 0.01487 |

We tested all the bottom-up (tree density, frequency and genetic susceptibility of chestnut tree) and top-down (hyperparasitism rates by native fungi and parasitoids) factors as fixed effects in the model before to sequentially removed non-significant parameters. While the GLM analysis including all the bottom-up factors in both sites showed significant p-values between 0.0331 and 0.0516, the best model was reached when removing the effect of genetic determinants. Indeed, the p-values obtained for the tree density and chestnut tree frequency in both sites are then lower and highlight that they are the main determinants in explaining the proportions of oscillatory dynamics among the predicted patterns.

**Supplementary Material 7. Variations in global biological control efficiency with respect to the dispersal of *D. kuriphilus* and *T. sinensis*.** Global rate of biological control was predicted for all combinations of sites as in SM5, with similar (A-C) or dissimilar (D-S) population dynamics predicted at the local scale, and while *D. kuriphilus* and *T. sinensis* rates of dispersal were varied from 0 to 50%. Colors indicate the variations in the predicted global control rate under the influence of *D. kuriphilus* and *T. sinensis* dispersal as represented in the legend. The black lines delineate the conditions where *D.kuriphilus* and *T. sinensis* co-exist with stable or oscilatory dynamics (E-G).

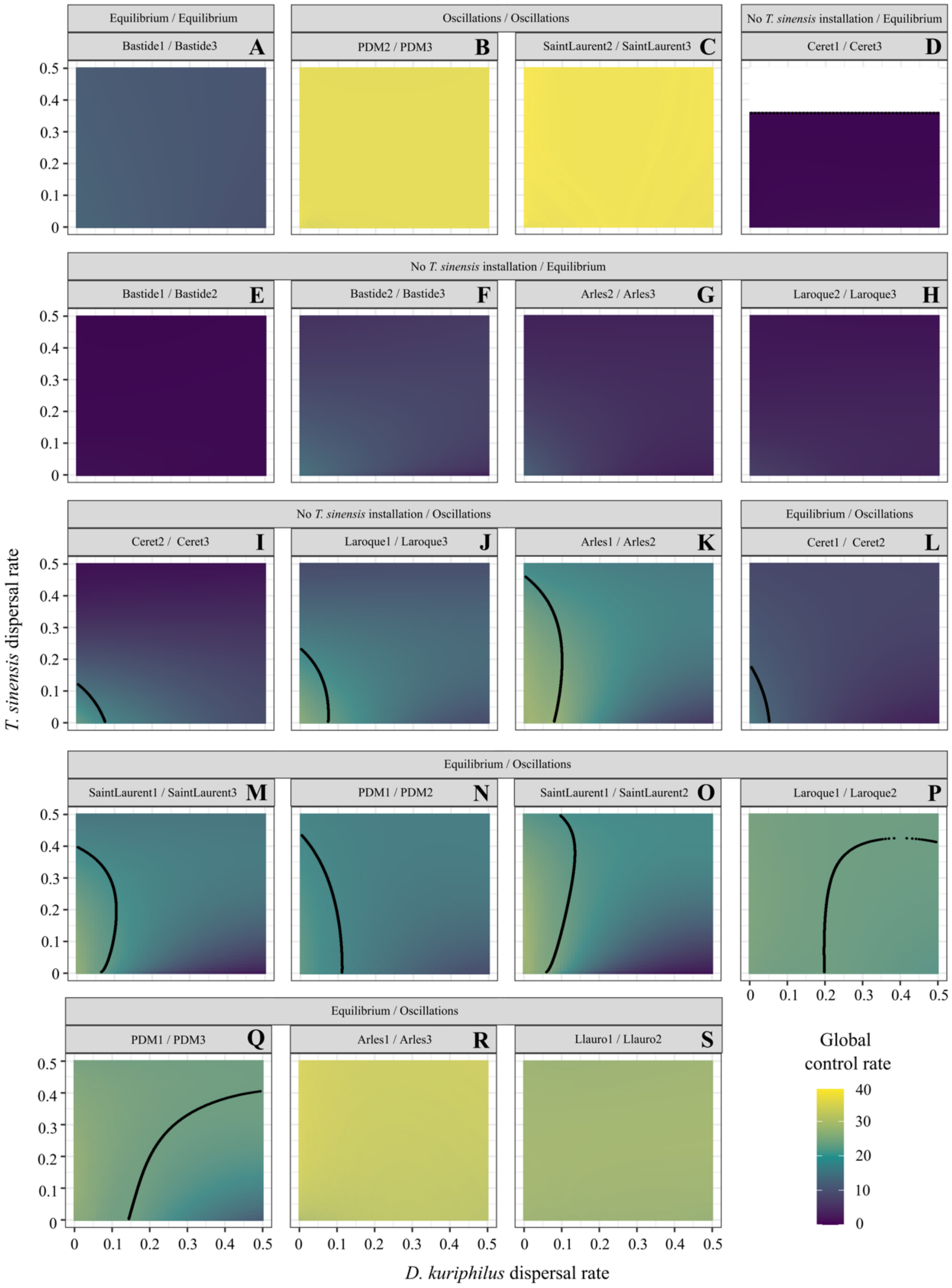

**Supplementary Material 8.** Variations in local biological control efficiencies with respect to the dispersal of *D. kuriphilus* and *T. sinensis*. Local rates of biological control were predicted for the same combinations of sites as in Figure 5, with similar (A-D) or dissimilar (E-P) population dynamics in the two connected sites, and while *D. kuriphilus* (x-axis) and *T. sinensis* (y-axis) rates of dispersal were varied from 0 to 50%. Colors indicate the variations in the predicted local control rates according to *D. kuriphilus* and *T. sinensis* dispersal rates as represented in the legend.

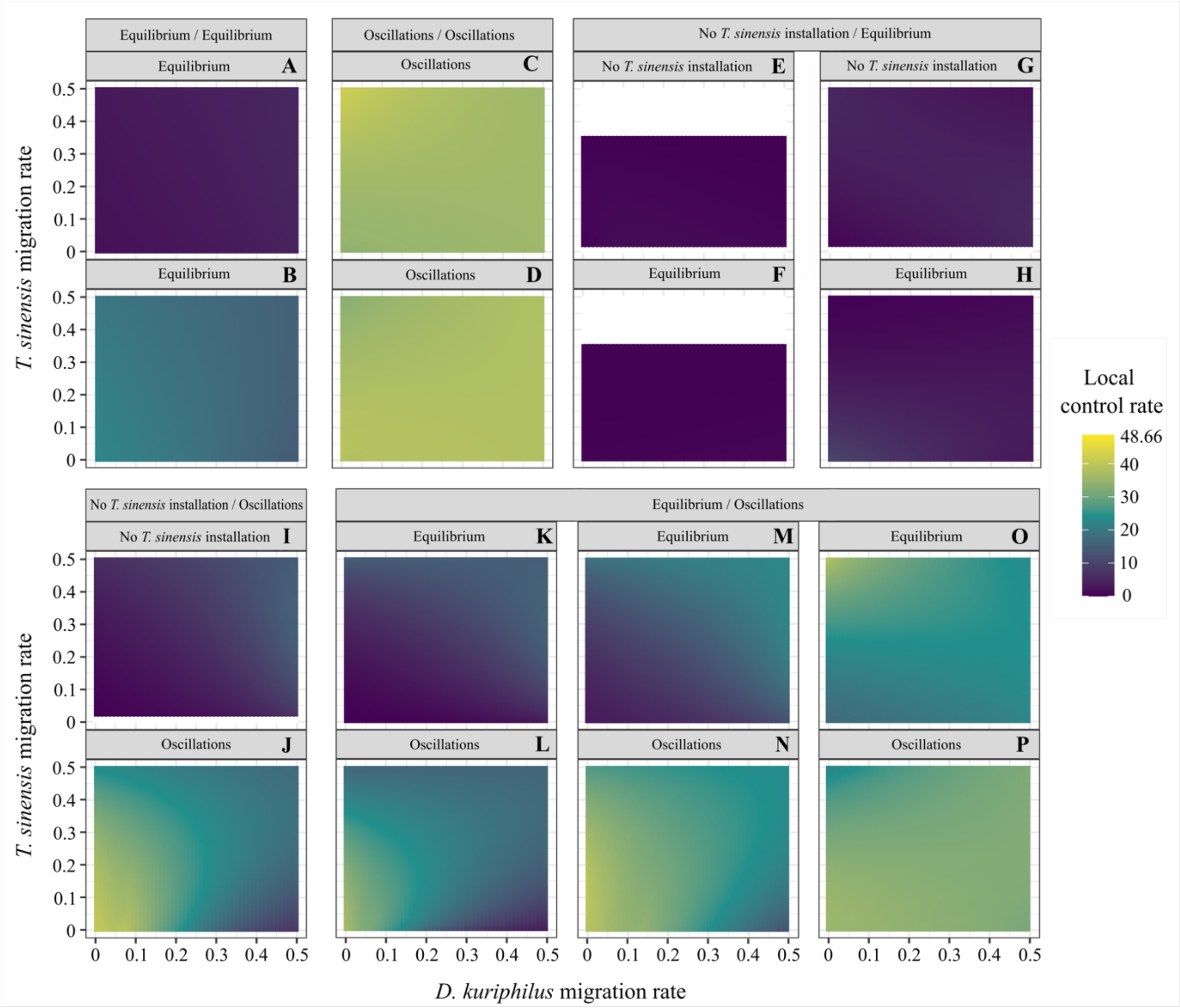
